# Membrane Curvature Generation by the Caveolin 8S Complex and the Role of Cholesterol

**DOI:** 10.64898/2026.02.11.705414

**Authors:** Sayyid Yobhel Vasquez Rodriguez, Themis Lazaridis

## Abstract

The protein caveolin-1 (cav1) is essential in the generation of caveolae, cup-like invaginations in the plasma membrane, but the mechanism of its action remains unclear. A recent cryo-EM structure revealed an 11-mer of cav1 (the 8S complex) forming a disk with a flat membrane-facing surface, raising the question of how a flat complex is able to generate membrane curvature. We previously conducted implicit-solvent and all-atom molecular dynamics simulations, which showed the 8S complex adopting a conical shape. The ability of the conical shape to remodel membranes was assumed but could not be confirmed. Here, we simulated the complex in discontinuous membrane patches of ∼30 nm diameter on the Anton 3 supercomputer. In a 2-μs simulation, a flat POPC membrane patch acquired pronounced positive curvature (curved away from the complex), converting into a hemisphere of 21-nm outer diameter. However, when the complex was constrained to prevent the conversion into a conical shape, the membrane patch acquired slightly negative curvature. These results show that the conical shape of the 8S complex is essential for positive curvature generation. A homology model of cav3 behaved very similarly to cav1, but the two recently discovered nonvertebrate caveolins remained flat and generated pronounced negative curvature. Simulations of cav1 in 70:30 POPC:cholesterol and other cholesterol-containing mixtures showed significantly lower curvature than in the pure POPC membrane or in an E. coli membrane mimic. This appears to be caused by cholesterol flipping from the distal to the proximal leaflet. No specific binding of cholesterol to the cav1 CRAC motif was observed, nor significant enrichment of cholesterol in contact with the complex. These observations lead to the hypothesis that cholesterol is enriched in caveolae not because of specific binding to caveolin, but because it can alleviate curvature stress due to its negative spontaneous curvature and its ability to rapidly flip-flop.

## INTRODUCTION

Caveolae are 50-100 nm wide invaginations of the plasma membrane of many mammalian cell types, which are enriched in cholesterol and sphingolipids. ^1^Their possible functions include resistance to mechanical stress, signaling, regulation of lipid homeostasis, transcytosis, and endocytosis ^2^. Two protein families are essential for the biogenesis of caveolae: caveolins and cavins ^3^. Caveolins appear to form the inner layer and cavins the outer layer of a protein coat that stabilizes caveolae ^4^. Mammals have three caveolin genes: caveolin-1 (cav1) is the best studied and most widely expressed, caveolin-3 occurs in muscle tissues, and caveolin-2 is inactive by itself but makes heterooligomers with cav1 ^5^. Membrane curvature in caveolae is assumed to arise from the presence of these protein components. cav1 alone can deform the plasma membrane when heterologously expressed in E. coli ^6^ and was shown to produce wide pan-like invaginations when cavin1 was knocked out ^7^.

Structural information on the caveolar protein coat is limited. cav1 has 178 amino acid residues and forms an 8S oligomer whose structure has been recently determined to be an 11-mer disk with a flat membrane-facing surface ^8^. The hydrophobicity profile of this disk suggested that it could be embedded deeply in a membrane displacing the proximal lipid leaflet. This raised the question of how a flat disk can bend the membrane. One possible mechanism was proposed based on the assumption that the lipid-protein interaction is less favorable than the lipid-lipid interaction, creating tilt and splay in the distal leaflet ^9^. Coarse-grained simulations also showed the flat disk creating small or large protrusions on a bilayer ^10,11^. All-atom simulations in POPC/cholesterol membranes showed only local curvature and suggested that the coarse-grained force field exaggerates membrane curvature ^12^.

We previously conducted molecular dynamics simulations of the cav1 8S complex in an implicit solvent membrane and observed it taking a conical shape with a concave membrane-binding surface and its outer ridge buried deep within the membrane ^13^. Subsequently, we simulated the CAV1 8S complex in all-atom POPC lipid bilayers and also observed the transition to a conical shape ^14^. This conical shape was inferred to be able to generate positive curvature, based on differences in binding energy to different size vesicles. However, this was not shown explicitly. The systems studied by all-atom simulations ^14^ were too small and too constrained by periodic boundaries to allow an assessment of the ability of the 8S complex to remodel membranes. The work of Liebl and Voth ^12^, which only showed local curvature, used a much larger POPC:cholesterol bilayer but was still constrained by periodic boundaries.

Here, to answer definitively the question of curvature generation we simulated the 8S complex in large, discontinuous membrane patches of different lipid compositions. The absence of any constraint reveals the intrinsic ability of the 8S complex to shape membranes. We also show the significance of the conical shape, by simulating the 8S constrained to remain flat as in the cryoEM structure. Simulations of nonvertebrate caveolins revealed a nonuniversal behavior of these complexes, and simulations with cholesterol led us to question past assumptions and propose a new hypothesis for the role of cholesterol in the biogenesis of caveolae.

## METHODS

All systems except the pure membrane started from the cryo-EM structure of the cav1 8S complex (PDB 7SC0) ^8^. First, continuous membranes with periodic boundary conditions were created in the CHARMM-GUI server ^15–17^. These systems were modified in the following ways: lipids beyond a circular or square patch of ∼27 nm diameter or edge were deleted and the place they occupied was filled with water. Because the C-terminal β-barrel is unstable without internal lipids, a few lipids were placed inside it (as a control, in one case the barrel was left empty, see SI). Several lipid compositions were simulated: pure POPC, 70:30 POPC:cholesterol, 65:25:10 POPC:cholesterol:PIP3, an E. coli inner membrane mimic (5:20:5 POPE:POPG:TOCL, where TOCL is tetraoleoyl cardiolipin), and an asymmetric mammalian membrane mimic (inner leaflet: 30% cholesterol, 10% POPC, 35% POPE, 25% POPS; outer leaflet: 30% cholesterol, 60% POPC, 10% POPE). The complex was inserted into the inner leaflet. For PIP3 a mixture of the three isomers (SAPI33, SAPI34, SAPI35, differing in which phosphate is protonated) was used. In all cases the complex was inserted deeply, displacing the proximal leaflet. The systems were equilibrated locally using NAMD following the standard six-step CHARMM-GUI protocol and then subjected to simulations on the Anton 3 supercomputer. All systems were run at 310 K (one system was rerun at 303 K to test possible effects of temperature, see SI). Each system contained over 2 million atoms. In one case the complex was restrained to remain close to the cryoEM structure using an RMSD restraint with force constant 65 kcal/mol/Å. The water thickness was set to 32 Å. The system was neutralized with potassium ions in addition to 0.15 M KCl. All systems used the TIP3P model for water ^18^ the charmm36m force field for proteins ^19^ and the charmm36 force field for lipids ^20^. As a control, one system was rerun with the Amber 19SB force field for proteins ^21^ and the Amber 21 force field for lipids ^22^(see SI). A few replicates were run to test reproducibility (see SI).

Simulation in DDM detergent started with building a 600-molecule micelle of BDDM (N-dodecyl-β-D-maltoside) molecules in the charmm-gui micelle builder ^23^ using the Martini 2.2 coarse grained force field ^24^. This was run with gromacs for 0.5 μs and then converted to atomistic representation using the charmm-gui all-atom converter. Sugar ring penetrations were resolved by deleting 28 DDM molecules and reequilibrating. This system was then run on Anton 3, first for 1 μs constraining the protein complex to remain close to the cryoEM structure, and then for another 2 μs releasing all constraints. This system contained the 8S complex, 572 DDM molecules, and 242,220 TIP3P molecules. A second system was created from the original one by adding 512 DDM molecules placed on a regular grid and deleting 123 that overlapped with existing DDM or protein. This system, with 961 DDM molecules, was equilibrated and subjected to the same Anton 3 protocol: 2 μs constraining the complex to remain close to the cryoEM structure, and 2 μs unconstrained.

Homology models for cav2 and cav3 were built using the swissmodel web server ^25^. The full-length model was constructed by aligning the AF2 prediction to each of the 11 protomers of 7SC0 and then adding the coordinates of residues 1 to 48. Details of all systems run are provided in Table S1 in SI.

## RESULTS

### A POPC membrane patch with the 8S complex acquires strong curvature

We started with a pure POPC membrane patch. As in previous work, the complex rapidly acquired a conical shape. Remarkably, the membrane patch takes the form of a spherical vesicle, open in the bottom and with the 8S complex on the outside (Fig. 2). The outer diameter of the sphere is about 21 nm. This happens rapidly; already at 200 ns there is pronounced curvature and after 0.5 μs the system is stable in the form shown in Fig. 2. The 7 POPC lipids in the barrel do not exchange in the 2 μs of the simulation. However, they are probably overcrowded; the barrel distorts substantially and a couple of lipids poke out through the β strands. Due to periodic boundary conditions the bottom lipids approach the complex from above and this prevents the closing of the vesicle. It is also noted that curvature exists even quite far from the complex. This may be due to line tension, the tendency of the open membrane patch to close to avoid the unfavorable exposure of hydrophobic acyl chains to water. However, it is clear that the initial bending of the membrane in a positive direction is caused by the complex (see also below). This picture is remarkably similar to that obtained using implicit membrane modeling ^13^.

**Fig. 1.**
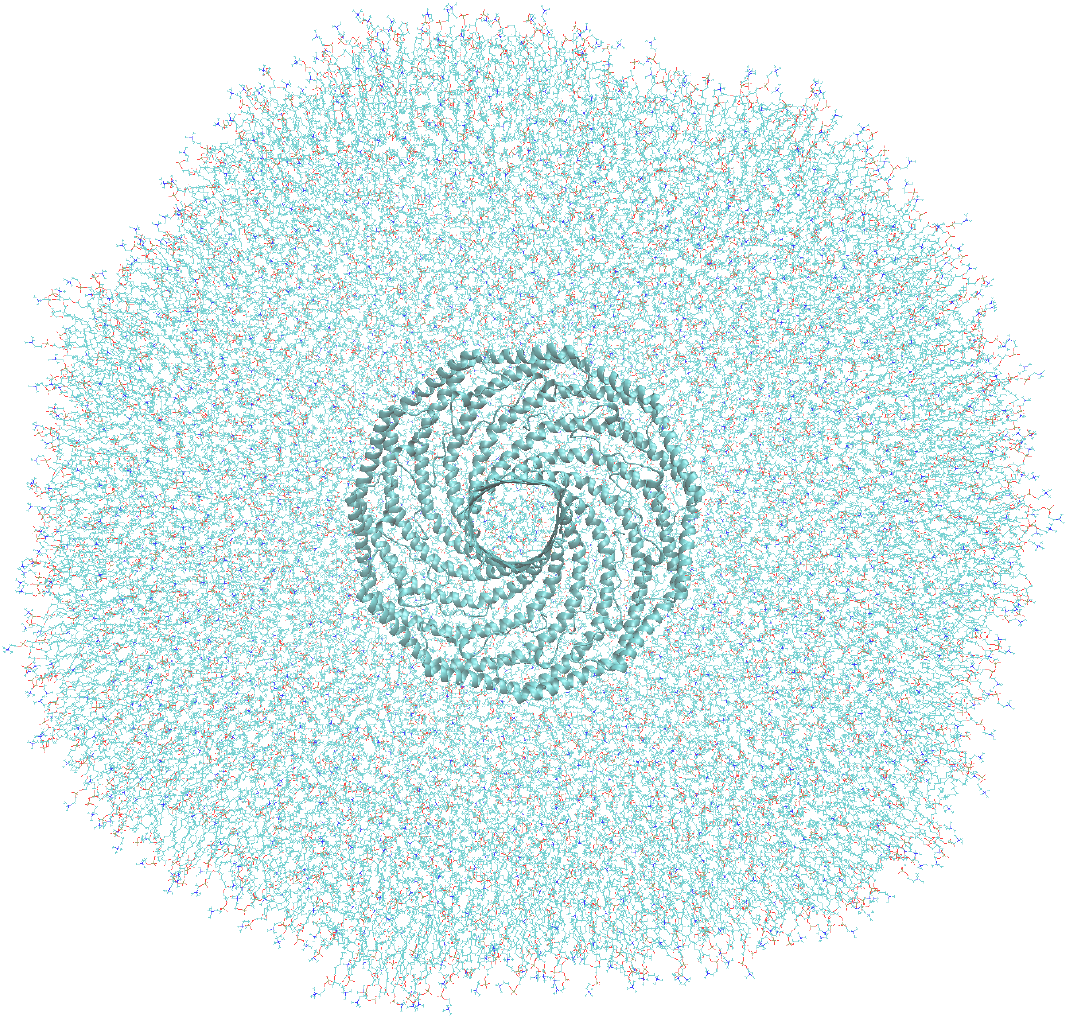
Top view of the starting configuration of the 8S complex in a circular membrane patch. The diameter of the complex is about 13 nm and that of the patch 31 nm. In some systems the patch was square instead of circular.

**Fig. 2.**
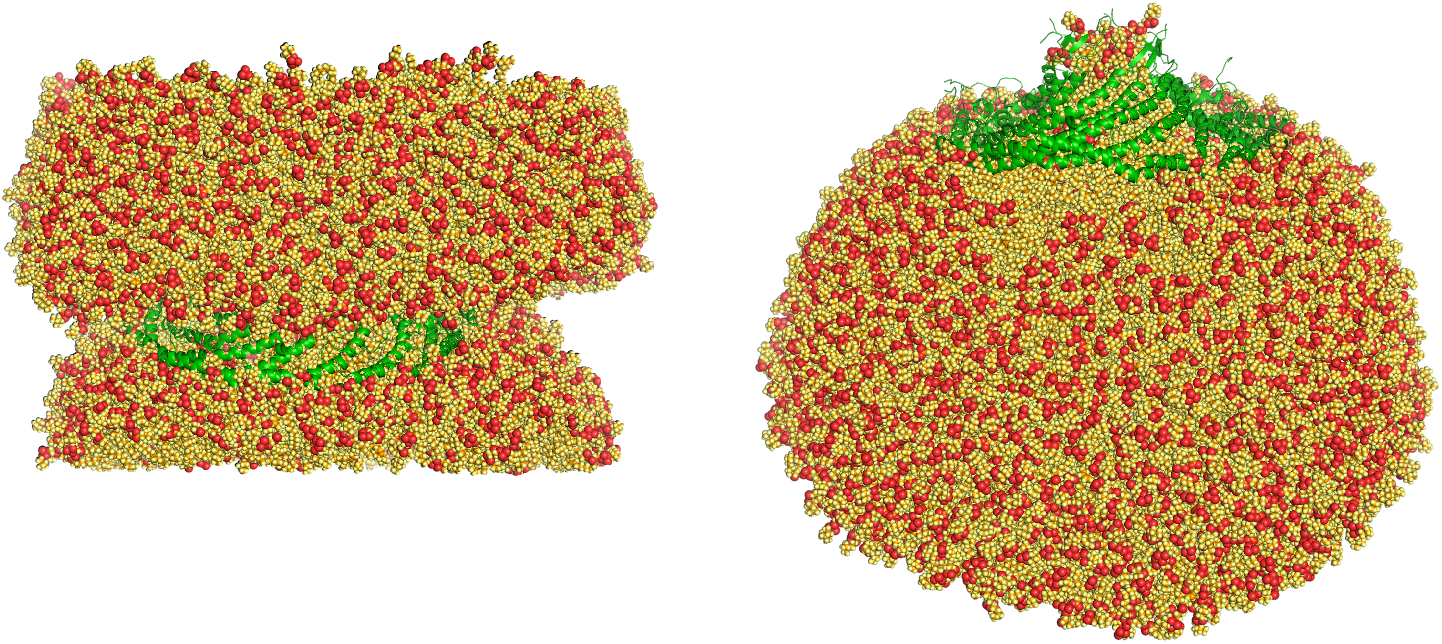
Final configuration of the POPC membrane with 8S. Left: original simulation cell. Right: the lipids have been wrapped to create a continuous membrane patch. The sphere is open at the bottom, as the lipids are interacting with image protein. The protein is in green ribbon. The red particles are P and O, orange particles are C, and all other lipid atoms are yellow. Water and ions are omitted.

### An 8S complex constrained flat generates slightly negative curvature

As in our previous work ^13^, in all simulations the cav1 8S complex becomes conical. To ascertain the role of the conical shape in membrane curvature generation, we ran the same simulation constraining the complex to remain close to the original, flat shape. Now the membrane exhibits slightly negative curvature (Fig. 3). This result shows that the conical shape is key to the generation of positive curvature.

**Fig. 3.**
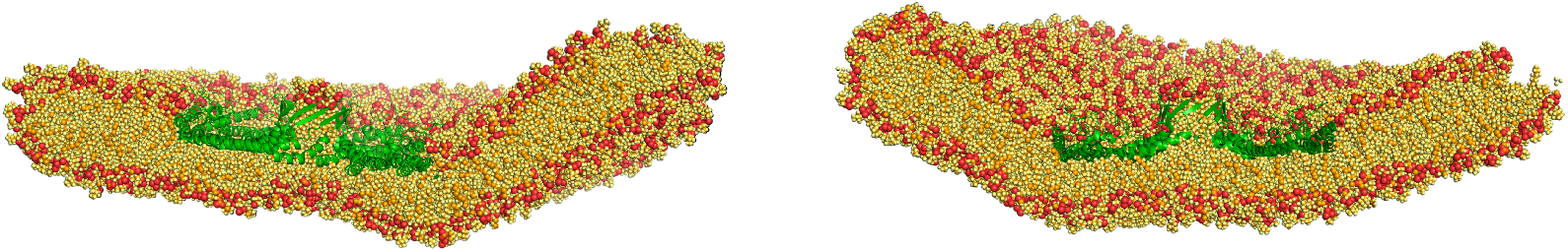
Final configurations of the POPC membrane with 8S constrained to remain close to the cryoEM structure. Two replicates are shown.

### Two nonvertebrate 8S complexes remain flat and generate negative curvature

Caveolins have been discovered in a wide range of organisms, including many that do not form caveolae ^26^. Recent work determined the structures of two 8S complexes from nonvertebrates, one from a sea urchin and one from a choanoflagellate ^26^. These structures are very similar to cav1 8S, except for a slightly more convex membrane binding surface. In implicit solvent runs (not shown) these structures remained flat. In the membrane patch all-atom simulations the structures also remained flat and, interestingly, created strongly negative membrane curvature, i.e. the complex was *inside* the vesicle formed (Fig. 4). This seems to be caused by interactions of the lipids with the rim of the complex (Fig. 5). The hydrophobicity of the rim and the presence of positively charged residues which can interact with lipid phosphates, draws lipids from the bilayer upward. Neighboring lipids follow them, allowing the bilayer to wrap around the complex. The outer diameter of the vesicles is about 20 nm.

**Fig. 4.**
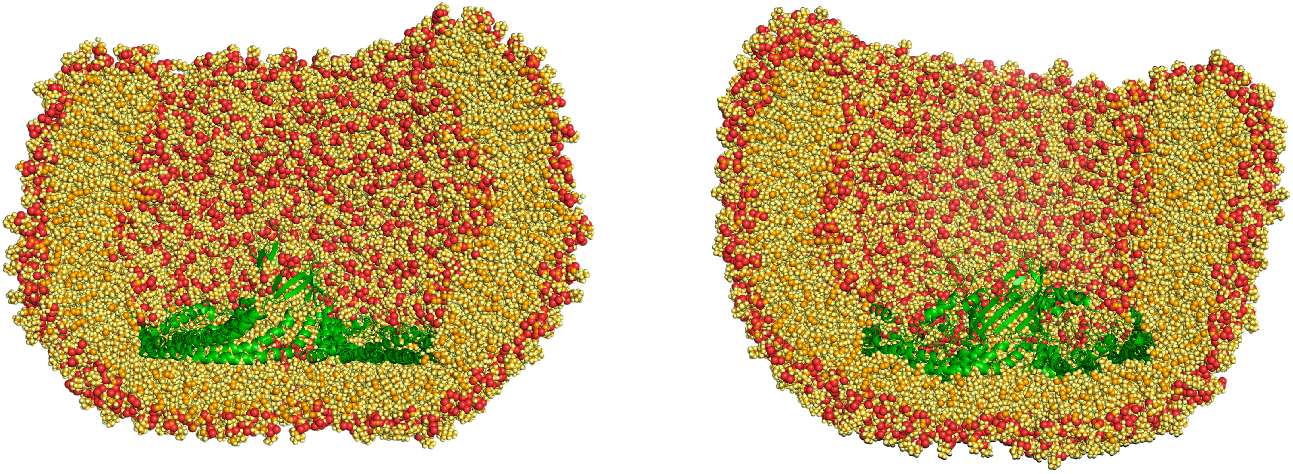
Nonvertebrate caveolin 8S after 2 μs of unconstrained MD simulation. A) 9DN0, S. purpuratus sea urchin B) 9DN1, S. rosetta choanoflagellate. The complex is inside the vesicle formed.

**Fig. 5.**
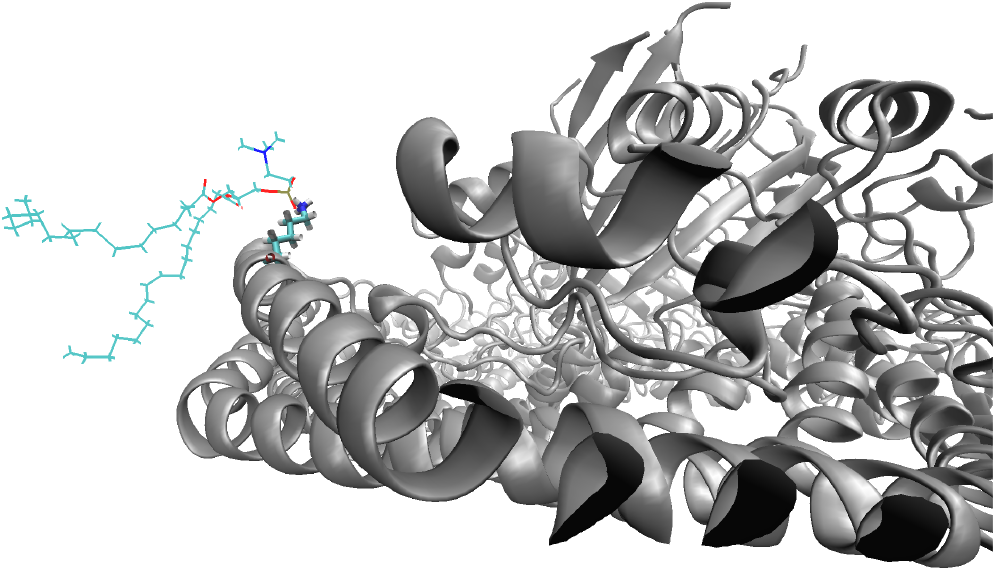
A POPC lipid at the rim of the 9DN1 8S complex. The lipid phosphate is interacting with a K132 residue.

The negative curvature generated by the flat cav1 complex is smaller than that generated by the nonvertebrate caveolins. This appears to be caused by the position of the positively charged residues in the rim. K86 in cav1 is lower and more outward oriented than the equivalent residues K59 and K132 in 9DN0 and 9DN1. Thus, POPC is forced to move higher up on the rim to interact with these residues, generating stronger curvature.

### Effects of lipid composition

Caveolae are known to be enriched in cholesterol ^27^ and cholesterol is required for the reconstitution of caveolin 1 into liposomes ^28^. It was thus of interest to consider the effects of cholesterol by simulating a 70:30 POPC:cholesterol bilayer. In addition, PIP3 was found to be enriched around cav1 ^29^, so we also considered a 65:25:10 POPC:chol:PIP3 membrane. The observation of heterologous caveolae in E. coli led us to consider an E. coli membrane mimic consisting of 75:20:5 POPE:POPG:TOCL. Finally, mammalian membranes are asymmetric, so we constructed an asymmetric mammalian membrane mimic composed of 30% cholesterol, 60% PC, 10% PE in the outer leaflet and 30% cholesterol, 10% PC, 35% PE, 25% PS in the inner leaflet, where the 8S complex resides.

The results are shown in Figs. 6-9. All systems created positive curvature, but the curvature was significantly smaller in systems containing cholesterol. The outer diameter of the curved membrane patch in 70:30 POPC:cholesterol was ∼ 50 nm (Fig. 6).

**Fig. 6.**
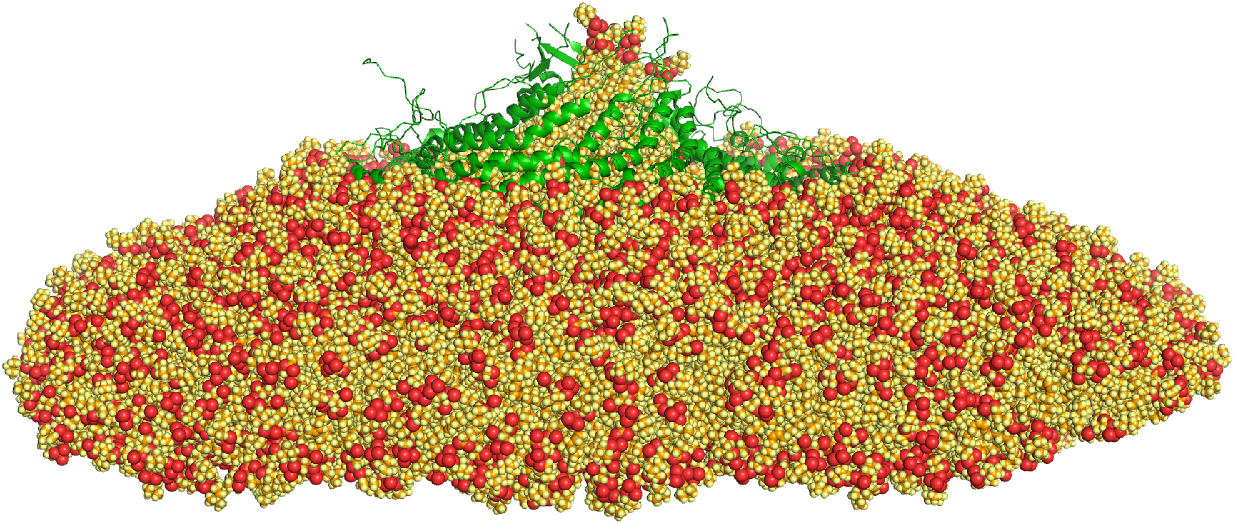
Cav1 8S in POPC-cholesterol after 4 μs of simulation.

The POPC-chol-PIP3 system gave similar curvature as the POPC-chol system (outer diameter ∼60nm) (Fig. 7). PIP3 molecules were seen to interact strongly with K86 and R101 in the scaffolding domain and a few R54 residues (residues 49-80 take different conformations in the 11 protomers. Some R54 residues remain far from the membrane and some reach out to interact with lipids.)

**Fig. 7.**
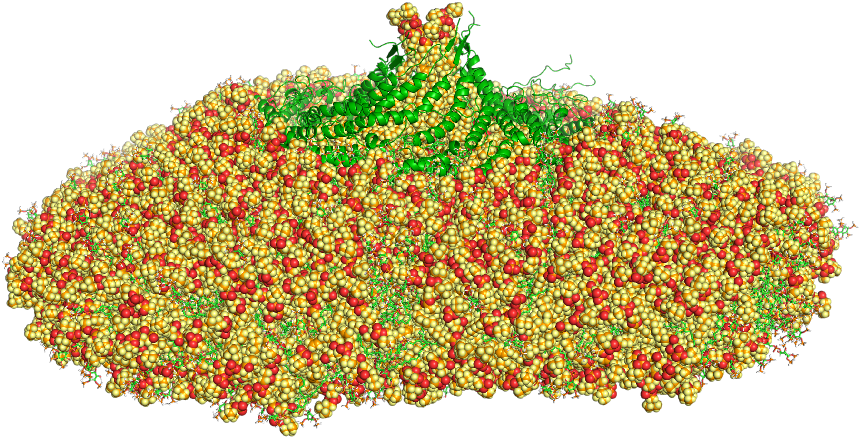
Cav1 8S in POPC-cholesterol-PIP3 after 2 μs of simulation. In green lines are the PIP3 molecules.

The E. coli mimetic gave large positive curvature (Fig. 8). The outer diameter of the membrane patch was about 21 nm, as in the pure POPC system, and somewhat smaller than the average of 30-33 nm found for h-caveolae ^30^. The negatively charged lipids POPG and cardiolipin are often seen to interact with K86 and R101. The different shape of the patch compared to POPC might be due to image interactions: the presence of negatively charged lipids in the E. coli mimetic which might prefer to interact with the rim of the complex in the neighboring cell.

**Fig. 8.**
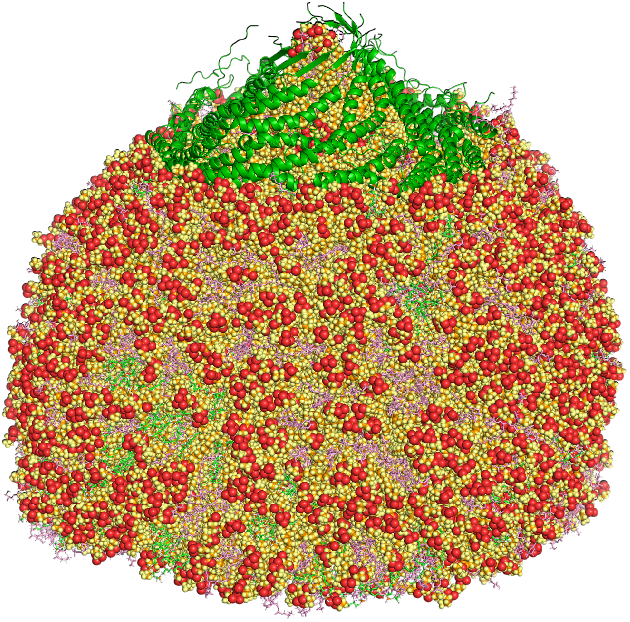
Cav1 8S complex in an E. coli mimetic membrane patch. Green represents TOCL and purple POPG.

Finally, the mammalian asymmetric model behaved similarly to other cholesterol-containing mixtures, generating low curvature (Fig. 9).

**Fig. 9.**
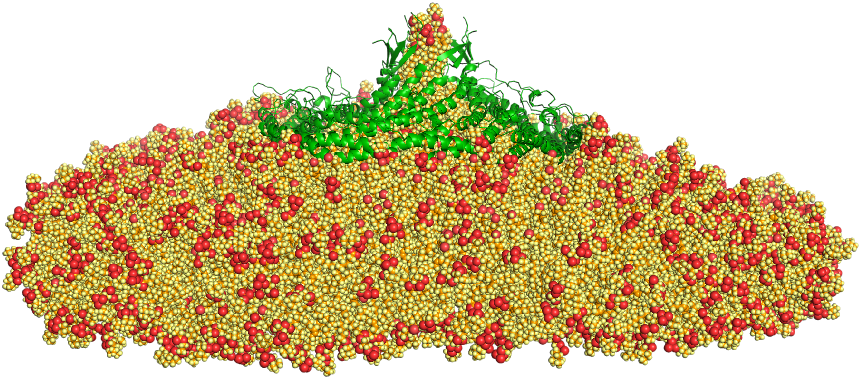
Cav1 8S complex in a mammalian asymmetric membrane after 1.14 μs of simulation.

### Analysis of the effect of cholesterol

We investigated the reasons for lower curvature in systems containing cholesterol, starting with POPC:cholesterol. Because cholesterol is known to have a negative intrinsic curvature (small headgroup, bulky tail) ^31^, we examined the distribution of cholesterol in the final configuration and observed a larger number of them in the proximal leaflet than at the beginning of the simulation. Specifically, in the final 4-μs structure we observe 74 flips (from distal to proximal leaflet) and 59 flops (from proximal to distal leaflet) compared to the initial structure. So, there was a net of 15 flips to the proximal leaflet. This reduces both the positive curvature of the proximal leaflet and the negative curvature of the distal leaflet, leaving the bilayer less curved. In the 2-μs replicate simulation the net number of flips was 7. The observed number of flip-flops is comparable in other recent simulations ^32,33^.

Caveolin contains a so called CRAC motif in the C-terminal region of the scaffolding domain, residues 94-101 (VTKYWFYR). The CRAC motifs, defined as L/V-X^1−5^-Y-X^1−5^-K/R, have been suggested as cholesterol binding elements ^34,35^, but their predictive value has been questioned ^36,37^. To check whether this motif has any special interaction with cholesterol, we calculated the interaction energy of residues 94-101 with either cholesterol or POPC. The POPC interaction energy was scaled by 3/7 to account for abundance and by 74/134 to account for the larger number of atoms in POPC vs. cholesterol. The results are plotted in Fig. 10. The interaction energy with cholesterol does not increase with time and is actually smaller in magnitude than that with POPC, despite the normalization. This is due in part to electrostatic interactions between the two basic residues (K86 and R101) and phosphate in POPC. In addition, we looked at the exchange of specific cholesterol molecules in any of the 11 CRAC sites. In most cases specific cholesterol molecules come in and out of contact with a CRAC motif quite rapidly. The largest residence time observed was 730 ns. These results do not support any special role of these sites in cholesterol recognition.

**Fig. 10.**
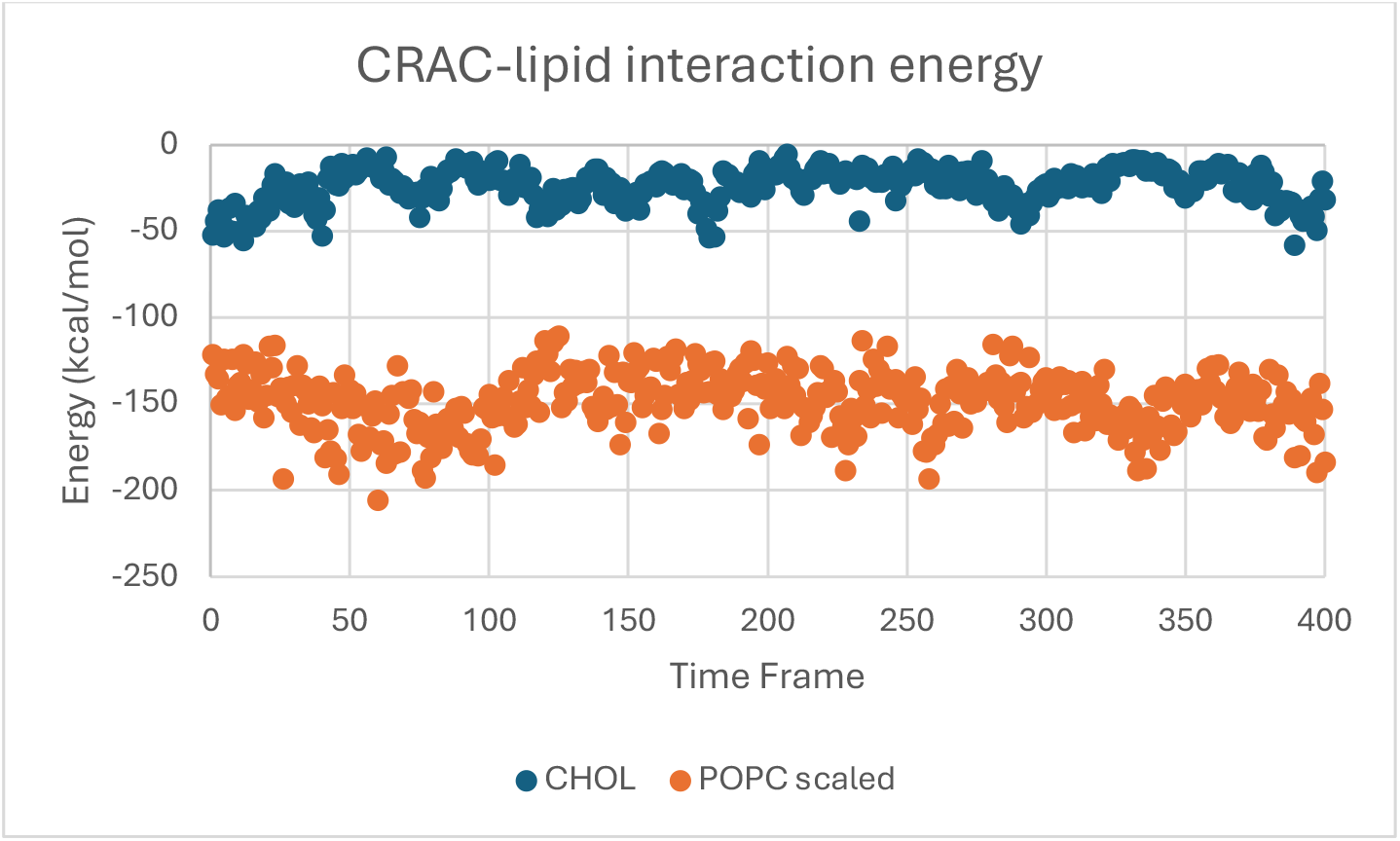
Interaction energies between the CRAC residues and either cholesterol or POPC. The values for POPC have been scaled by 3/7 × 74/134 (see text). Each time frame corresponds to 10 ns. Similar results were obtained for the CARC motif (residues 96-103, not shown).

We also checked whether cholesterol is enriched around the complex. For that, we calculated the number of cholesterol molecules within 15 Å from any protein atom as a function of time (Fig. 11). With an initial drop and subsequent rise and drop, this number is essentially constant throughout the simulation.

**Fig. 11.**
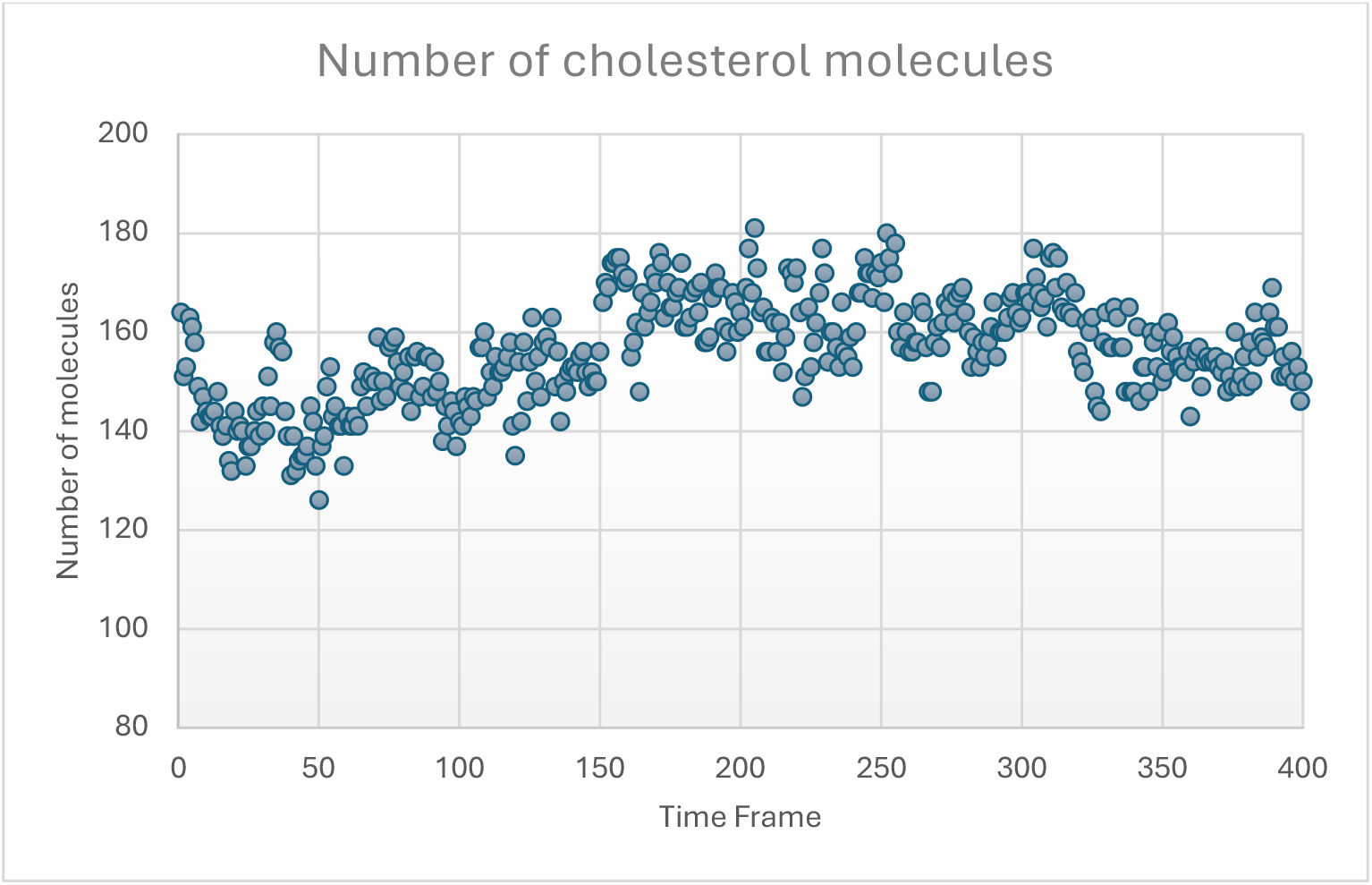
Number of cholesterol molecules within 15 Å from any protein atom as a function of simulation frame (each frame corresponds to 10 ns).

Fig. 12 shows a snapshot of the cholesterol hydroxyl oxygen atoms in the final frame of the POPC-cholesterol simulation. The protein complex (not shown) is near the center of the circle. Again, no increased cholesterol density is observed under the complex.

**Fig. 12.**
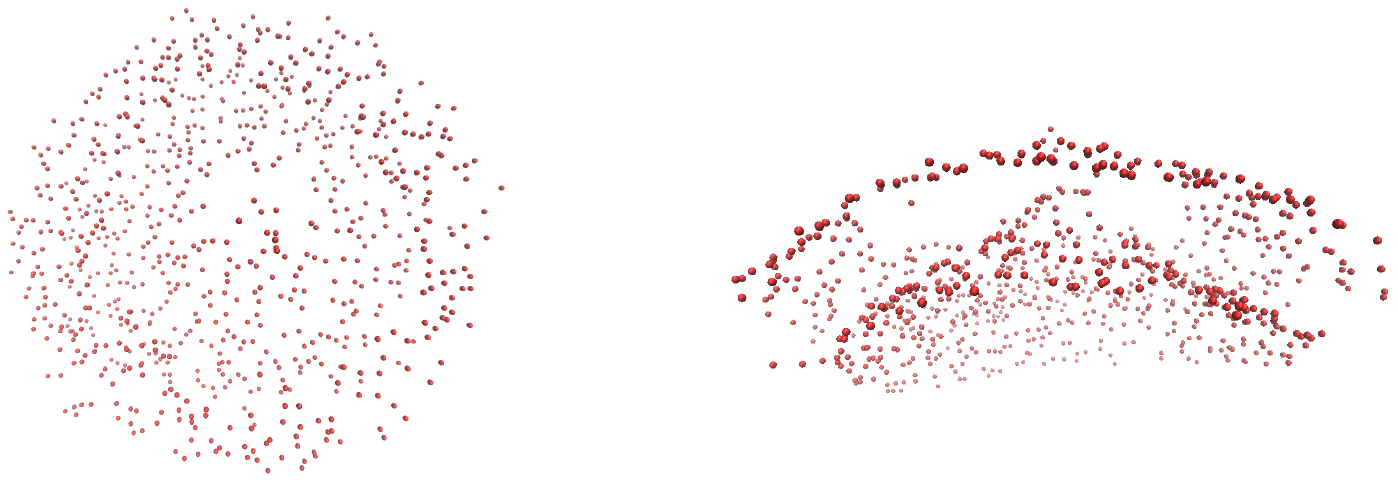
Distribution of cholesterol O3 atoms in the last frame of the 4-μs simulation in POPC-cholesterol. Top view (left) and side view (right).

One might ask whether 4 μs is enough time for the lipids to diffuse into or out of contact with the complex. So, we calculated the rmsd for each lipid between the initial and final configuration relative to the complex. This value was 106 Å for cholesterol and 120 Å for POPC. The longest distance of a lipid from the complex in the beginning is about 114 Å. Thus, the lipids are mobile enough to show enrichment or depletion in the timescale of the simulation, if that is thermodynamically favored.

In the POPC:cholesterol:PIP3 system analysis of flip-flops showed that a net of 4 cholesterol molecules flipped from the distal to the proximal leaflet, much fewer than in the POPC-cholesterol system. Similar analysis showed that PIP3 also showed a net of 7 flips to the proximal leaflet (it can do so over the open edge of the membrane patch). The spontaneous curvature of PIP3 is not known experimentally, but PIP2 prefers negative curvature ^38–40^. PIP3 appears to share this property and to contribute to lowering the curvature of this membrane patch together with cholesterol.

Analysis of flip-flops in the mammalian asymmetric model reveals a more complex picture. First, we should note that any open membrane patch will not remain asymmetric for very long, because any lipid can change leaflets over the open edge. Indeed, we see considerable movement of lipids between the two leaflets in the 1.14 μs of simulation. Unlike other systems, cholesterol here exhibits a net of 26 *flops* to the distal leaflet. POPE, which also has a negative spontaneous curvature ^41^, has an even larger number of net 68 flops. But we start with a lot more POPE in the proximal leaflet (585 proximal vs. 437 distal), so the proximal leaflet already has a negative curvature disposition, which it relieves by moving POPE to the distal leaflet. Several POPS molecules also move to the distal leaflet. These complications necessitate revisiting this model under conditions where the flip-flop of phospholipids will not be allowed. However, the observations aboe suggest that the presence of more negative-curvature lipids in the cytoplasmic leaflet of mammalian membranes could also play a role in alleviating the impact of positive curvature generating objects like the cav1 8S.

### Other caveolins

We constructed homology models of cav3 and cav2 based on the CAV1 8S complex structure. Cav3 behaved very similarly to cav1 (Fig. 13). Cav2 is experimentally known to not form 8S complexes ^42^. Nevertheless, on a 2-μs timescale it remained stable and generated negative curvature (see SI).

**Fig. 13.**
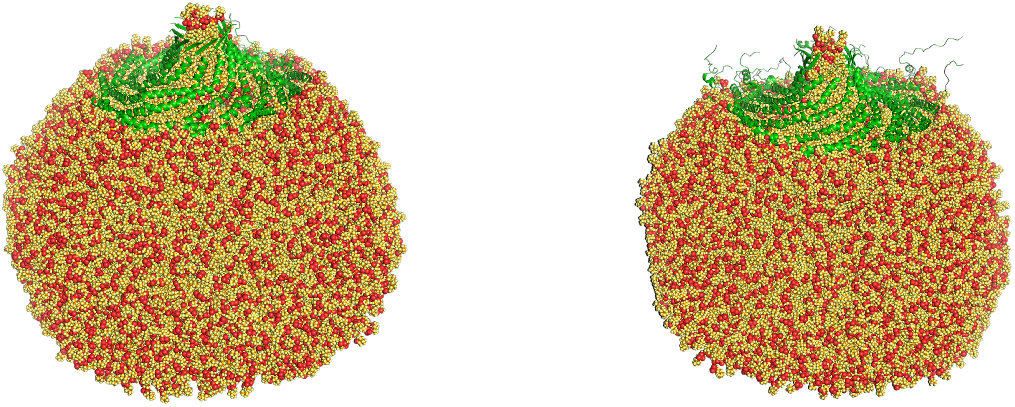
cav3 8S complex in POPC. Left: swissmodel, right: AF2 model

### Controls and variant runs

Several controls were performed to confirm the reliability of the results and are shown in the SI. First, a pure POPC membrane patch was run without protein for 2 μs. The patch remained flat for about 1.2 μs and then started curving. The final curvature is similar to that in the POPC/chol system, but in the latter it develops much faster, within 200 ns.

To test the effect of the force field we ran a simulation using the Amber 19SB force field for the protein and the Lipid21 force field for the lipids. The result was very similar to the charmm36 ff. We ran a simulation at lower temperature of 303 K instead of 310 K, and the result was again similar. Most runs were done with a semi-isotropic barostat. A couple of runs with an isotropic barostat gave very similar results.

Replicate simulations were done for a) the constrained 7SC0 complex in POPC, b) the POPC/chol system, c) the two nonvertebrate caveolins. The results were very similar (see SI).

Cav1 has three Cys that may be palmitoylated: 133,143,156. We simulated a system with 133 and 156 palmitoylated in POPC. This system behaved very similarly to the unpalmitoylated one (see SI). It developed a curvature of 1/11nm (outer diameter 22 nm). Some palmitoyl chains were extended and others ran parallel to the protein surface.

The first 48 residues are not visible in the cryoEM structure and are predicted to be disordered. However, they harbor functionally important residues, such as Y14 which can be phosphorylated ^43^. The 59 N-terminal residues were found important for generation of the higher 70S oligomer^44^, and thus are expected to be involved in interactions between 8S complexes. Here we used a (low confidence) AF2 prediction, which places the N-terminal region on top of the disk and predicts a helix at residues 28-42 and a β-hairpin at residues 4-15. This system exhibited the conical transformation and generated positive curvature similar to the truncated system (see SI).

### Simulations in DDM detergent

Although in all our simulations the cav1 8S complex converts to conical, the cryoEM structure is clearly flat, which is of significant concern. CryoEM structure determination was done in DDM detergent ^8^. To check whether that environment favors a flat 8S complex, we carried out two simulations in DDM, one with 572 and one with 961 detergent molecules. The first simulation started from a coarse-grained micelle, converted to all atom representation. In the second simulation additional DDM molecules were added in the solution surrounding the complex to check for possible effects of DDM concentration. In both simulations the complex was first constrained to the cryoEM structure and then the constraints were released. In both cases we observed the transition to conical conformation. In the second simulation most of the additional detergent formed a micelle next to the complex (Fig. 14). We conclude that DDM is not responsible for the discrepancy between simulated and cryoEM structures.

**Fig. 14.**
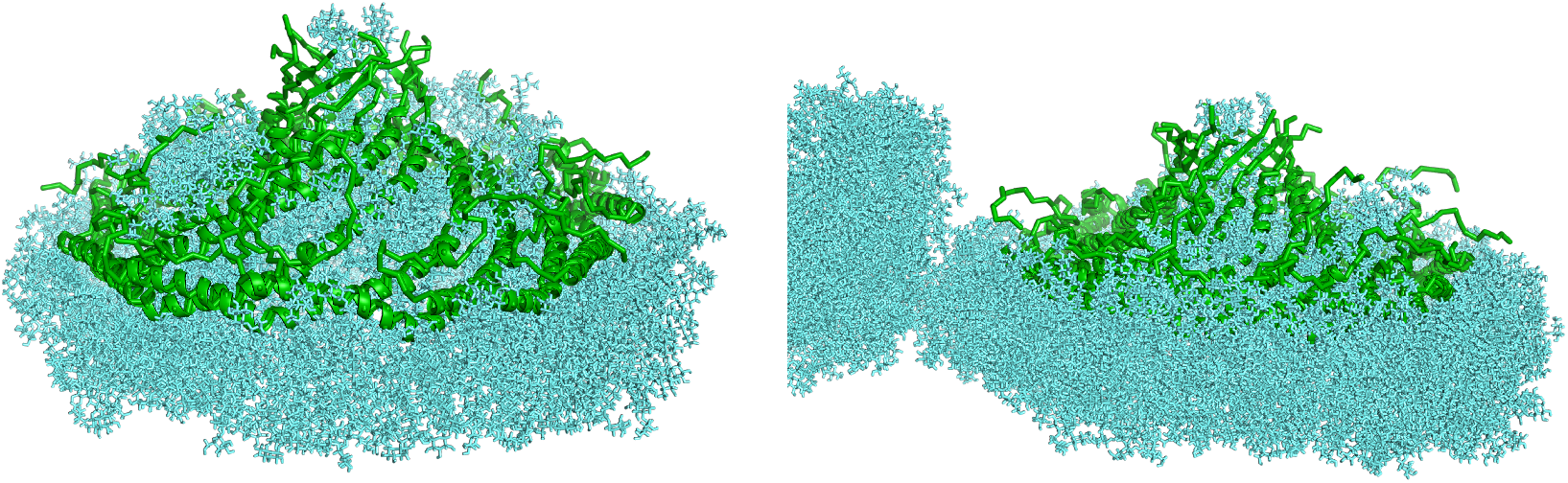
CAV1 8S complex in DDM detergent. A) 572 DDM molecules, b) 961 DDM molecules.

## DISCUSSION

This work shows that the cav1 8S complex has an intrinsic ability to bend membranes into spherical shape, if and only if it attains a conical conformation. This conclusion was reached by simulating large discontinuous membrane patches. This approach, pioneered by the Schulten group ^45^, is not without limitations. First, the large rearrangement of the membrane patch brings image lipids into contact with the central part of the patch and the top of the protein complex, affecting the final attained conformation. However, the direction of membrane bending is clearly not affected by this artifact. Second, the free membrane edge has an unfavorable free energy (line tension) that would by itself force the membrane to eventually close. So, any attempt to extract quantitative estimates of the bending force should account for the line tension. The line tension could be reduced in future work by adding some detergent at the free edge. Finally, real membranes in vivo or in vitro are not free patches but form closed surfaces. Whether a closed membrane will bend or not will depend on the tension and the availability of extra lipid to accommodate the increase in surface area.

It is interesting that different caveolin 8S complexes, despite a very similar structure, behave differently in simulations. The two nonvertebrate 8S complexes do not exhibit the transition to a conical conformation. This is consistent with the fact that they do not form heterologous caveolae in E. coli ^46^. The function of caveolin in these species is unknown. The amount of membrane bending appears to depend on two factors: the shape of the complex, and the distribution of hydrophobic and charged residues on the rim of the complex.

There is a lot of experimental evidence that cholesterol is crucial for caveolae formation. For example, cholesterol depletion flattens caveolae ^47–51^. Cholesterol oxidation leads cav1 to move from the plasma membrane to Golgi ^52^. Cav1 was successfully reconstituted into liposomes only if the cholesterol content was >30% and copurification assays led to the conclusion that Cav1 binds cholesterol ^28^. Caveolae are enriched in cholesterol & sphingolipids ^27^. Cholesterol is required for oligomerization of CAV1 into the larger 70S complex ^44^ and promotes caveola scission ^53^.

It is impossible to rationalize all these observations based on our limited data, but significant insights have been obtained. First, we do not observe any strong, specific binding of cholesterol to either the CRAC motif or to other parts of the protein. Previous computational work also cast doubt on the predictive ability of the CRAC motif ^37,54^. Here, by “binding” we mean long-lasting direct contacts. Small peptides that included the CRAC motif were found to promote cholesterol segregation ^55,56^, but peptides may have different conformation and interact differently with lipids than the same sequence in the context of a larger structure. Our result is in apparent contradiction to previous work that concluded that cav1 binds cholesterol ^28^. That conclusion was based on the fact that some cholesterol co-purified with caveolin. However, the conditions in that assay are far from native and the protein under those conditions is unlikely to be in the oligomeric form considered here. For many experimental techniques binding means enrichment in the broad vicinity of the protein, not necessarily a contact interaction. We note that our analysis was done on the unpalmitoylated protein and palmitate may interact favorably with cholesterol ^57^.

What we do observe is an ability of cholesterol to alleviate the curvature stress imposed by the complex on the bilayer due to its negative intrinsic curvature and its ability to flip-flop. This may explain the cholesterol requirement for cav1 insertion into preformed liposomes. A pure POPC liposome might not be able to accommodate the strong curvature imposed by the complex. Cholesterol allows protein incorporation with a much milder overall curvature. Thus, cholesterol enrichment in caveolae may be due not to specific interactions but due to the fact that cholesterol is needed in the vicinity of the complex to alleviate curvature stress.

The E. coli mimetic membrane patch bends as much as pure POPC. This seems consistent with the observation of caveola-like vesicles when cav1 is heterologously expressed in E. coli ^6^. The much lower curvature observed in the mammalian mimetic POPC/chol mixture explains why cav1 alone is sufficient for caveolae formation in E. coli but not in mammalian cells. Cholesterol in the latter makes it more difficult to form caveolae, thus offering more opportunities for regulation and dynamics and necessitating additional proteins, the cavins. These results are consistent with the fact that caveolin alone generates lower curvature in mammalian cells ^7^ than in prokaryotic cells ^30^.

Harder to explain is why cholesterol depletion flattens caveolae. If our hypothesis is correct, removing cholesterol should enhance curvature, not diminish it. However, the in vivo situation is quite complex. First, cholesterol depletion is always partial. Cyclodextrin may diminish cholesterol concentration by 50% ^48,49^ but substantial amounts remain that may be enough to alleviate curvature stress. Secondly, caveolae formation in mammals also requires cavins. It has been reported that cholesterol depletion causes degradation of cavin 2 and movement of cavin 1 to the cytoplasm ^51^. So, partial depletion of cholesterol may remove cavins, which are essential for caveolae formation, and at the same time leave enough cholesterol in the membrane to prevent budding. Furthermore, caveolae formation is more complex than simply creating spherical curvature. Caveolae have a neck region with negative Gaussian curvature which may have special lipid requirements ^58^. Cholesterol is known to facilitate membrane fusion, which requires the transient formation of necks ^59^. The lipid distribution between bulb and neck regions is unknown.

Two recent articles reported coarse-grained simulations of the 8S complex. Martini2 ^10^produced modest protrusions at the site of 8S binding, while Martini3 produced dramatic remodeling of a flat bilayer but not of a liposome ^11^. In both studies the 8S complex is essentially constrained flat by the coarse-graining process. Under this condition, our all-atom simulations produced negative curvature. Thus, there is a discrepancy between all-atom and coarse-grained simulations that should be analyzed and understood. Maybe the coarse-grained force fields cannot reproduce configurations like the one depicted in Fig. 5. The Martini2 study also found cholesterol enrichment under the complex, but we are not able to confirm the same in all-atom simulations. Liebl & Voth provided evidence that Martini2 exaggerates membrane curvature ^12^. They did not observe global curvature in their periodicically continuous POPC/chol bilayer. This observation is consistent with our finding that the POPC/chol system generates mild curvature. It would be interesting to repeat that simulation in pure POPC.

The observation of negative curvature for the flat complex also contradicts a model of 8S curvature generation based on lipid tilt and splay ^60^. These authors considered the complex to be a flat disk and assumed that lipid acyl chains in the distal leaflet will have lower affinity for the caveolin disk than for other lipids, so they will tend to avoid contact with it. This will create tilt and splay that will propagate and bend the membrane away from the complex. This was not observed in our simulations of the flat-constrained 8S complex. Instead, we observed slightly negative curvature due to lipid interactions with the rim of the complex, which are ignored in the lipid splay model.

A troublesome concern is that the cryoEM structure of CAV1 8S is undeniably flat. Detergent is known to sometimes stabilize structures distinct from those in bilayers ^61–63^. However, our simulations in DDM also show the transition to conical shape. In our hands the 8S complex remains flat only when simulated in vacuum ^14^. Here we also carried out simulations in explicit n-hexane and found distortions due to strong electrostatic interactions, but the barrel did not rise as in the other simulations (see SI). Water seems to weaken interactions in the spoke region and allow the shape to change to conical. Could it be that traces of organic solvent in DDM displace this water and keep the structure flat? Even 1% impurities could provide substantial numbers of small organic molecules to exert a visible effect. Alternatively, could the very low temperature of the cryoEM experiment somehow prevent water from exerting its “lubricating” function? Ultimately, experiments in native bilayers at physiological temperature will provide a definitive answer.

## Supporting Information

One table and 10 additional figures.

A zip file with the final coordinates of the key all-atom simulations in PDB format will be provided upon publication in a peer-reviewed journal.

## Acknowledgements

This work was supported by the National Science Foundation (MCB 1855942). Anton computer time was provided by the Pittsburgh Supercomputing Center through grant R01GM116961 from the NIH. The Anton machine at PSC was generously made available by D.E. Shaw Research, and computer time was provided by the National Center for Multiscale Modeling of Biological Systems through grant number P41GM103712-S1 from the NIH and Pittsburgh Supercomputing Center.

