## Supporting Information for "Membrane Curvature Generation by the Caveolin 8S Complex and the Role of Cholesterol"

Graduate Programs in Chemistry, Biochemistry, and Physics,
The Graduate Center, City University of New York,365 Fifth Ave., New York, NY 10016, USA

**Table S1**. Details of the systems run.

| | Atoms Total | Lipids Total | Proximal | Distal | Lipid in Barrel | TIP3P | K+/Cl- | Duration (µs) |
|---|---|---|---|---|---|---|---|---|
| POPC | 2429020 | 2207 | 1000 | 1200 | 7 | 702287 | 1567 / 1534 | 2 |
| Amber FF POPC | 2429020 | 2207 | 1000 | 1200 | 7 | 702287 | 1567 / 1534 | 2 |
| 303 K POPC | 2429020 | 2207 | 1000 | 1200 | 7 | 702287 | 1567 / 1534 | 2 |
| ISO POPC | 2429020 | 2207 | 1000 | 1200 | 7 | 702287 | 1567 / 1534 | 1 |
| POPC Run2 | 2429020 | 2207 | 1000 | 1200 | 7 | 702287 | 1567 / 1534 | 2 |
| CONS POPC | 2429020 | 2207 | 1000 | 1200 | 7 | 702287 | 1567 / 1534 | 2 |
| CONS POPC Run2 | 2429020 | 2207 | 1000 | 1200 | 7 | 702287 | 1567 / 1534 | 2 |
| POPC/CHOL | 2195764 | 1827 / 790 | 831 / 360 | 992 / 426 | 4 / 4 | 622131 | 1403 / 1370 | 4 |
| POPC/CHOL Run2 | 2195764 | 1827 / 790 | 831 / 360 | 992 / 426 | 4 / 4 | 622131 | 1403 / 1370 | 2 |
| POPC/CHOL no barrel lipids | 2194932 | 1823 / 786 | 831 / 360 | 992 / 426 | N/A | 622131 | 1403 / 1370 | 2 |
| POPC/CHOL ISO after 4µs SEMIISO | 2195764 | 1827 / 790 | 831 / 360 | 992 / 426 | 4 / 4 | 622131 | 1403 / 1370 | 2 |
| 9DN0 | 2508673 | 2205 | 1000 | 1200 | 5 | 729273 | 1571 / 1604 | 1.93 |
| 9DNO ISO Run2 | 2508673 | 2205 | 1000 | 1200 | 5 | 729273 | 1571 / 1604 | 1 |
| 9DN1 | 2725938 | 2205 | 1000 | 1200 | 5 | 800031 | 1729 / 1718 | 2 |
| 9DN1 Run2 | 2725938 | 2205 | 1000 | 1200 | 5 | 800031 | 1729 / 1718 | 1 |
| POPC/CHOL/PIP (SAPI 33-35) | 2166161 | 1617 / 688 / 85 / 94 / 85 | 724 / 324 / 41 /46 /38 | 888 / 363 / 44 / 48 / 47 | 5 / 1 / 0 / 0 / 0 | 610178 | 2971 / 1354 | 2 |
| POPE/POPG/TOCL2 (E. coli mimic) | 2322850 | 1817 / 501 / 123 | 839 / 227 / 50 | 973 / 272 / 73 | 5 / 2 / 0 | 658216 | 2203 / 1423 | 1.99 |
| POPC/CHOL/POPE/POPS (Mamm. Asymmetric) | 2055790 | 1015 / 839 / 585 / 312 | 115 / 395 / 437 / 312 | 896 / 442 / 147 / 0 | 4 / 2 / 1 / 0 | 572892 | 1647 / 1302 | 1.14 |
| Palmitoylated | 2447391 | 2205 | 1000 | 1200 | 5 | 708146 | 1581 / 1548 | 2 |

| | | | | | | | | |
|---|---|---|---|---|---|---|---|---|
| Cav1 full length | 2935652 | 2206 | 1000 | 1200 | 6 | 868134 | 1994 / 1939 | 2 |
| Cav3 swissmodel | 2482524 | 2208 | 1000 | 1200 | 8 | 720200 | 1531 / 1531 | 2 |
| Cav3 Alphafold | 2768635 | 2207 | 1000 | 1200 | 7 | 814174 | 1806 / 1762 | 2 |
| Cav2 swissmodel | 2098101 | 2152 | 1000 | 1147 | 5 | 594938 | 1256 / 1245 | 2 |
| POPC membrane only | 2402186 | 2000 | 1000 | 1000 | N/A | 710428 | 1451 / 1451 | 2 |
| P132L | 2463779 | 2207 | 1000 | 1200 | 7 | 713855 | 1567 / 1534 | 1.81 |
| S80E | 2481273 | 2206 | 1000 | 1200 | 6 | 719731 | 1578 / 1534 | 1.97 |
| DDM 572 | 797675 | N/A | N/A | N/A | N/A | 242220 | 0 / 665 | 1cons / 2 uncons |
| DDM 961 | 757927 | N/A | N/A | N/A | N/A | 218505 | 0 / 609 | 2cons / 2 uncons |
| Hexane | 306313 | N/A | N/A | N/A | N/A | 0 | 0 / 0 | 1.93 |

### Supporting text

A number of replicate runs and controls were run to ensure the reliability of the results.

a) A pure POPC membrane patch was run without the 8S complex. The patch remained flat for 1.3 µs, and then acquired curvature. The final curvature was R ~20 nm, similar to the POPC-cholesterol system.

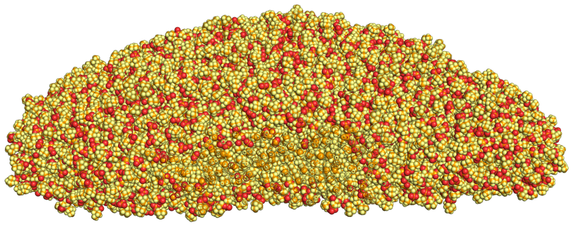

**Fig. S1**. POPC membrane without protein after 2 µs.

b) Replicate runs were conducted for the nonvertebrate caveolins, starting from different initial velocities. The replicates gave very similar results.

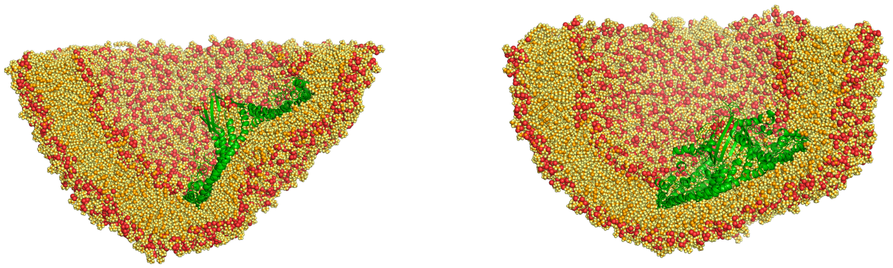

**Fig. S2**. Replicate runs of the nonvertebrate caveolin 8S complexes, 9DNO left and 9DN1 right.

c) In all runs lipids were added inside the β-barrel to stabilize it. To check the possibility that this may have affected the results, we ran the POPC/cholesterol system without added lipids. This system produced results indistinguishable from the original run. One cholesterol molecule entered the barrel.

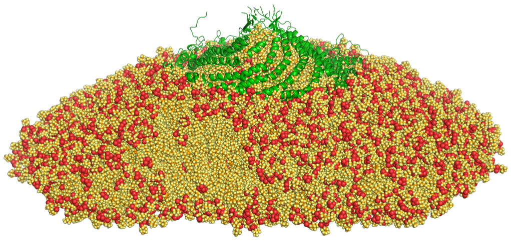

**Fig. S3**. 8S in POPC/cholesterol without added lipids in the barrel.

d) All simulations were run at 310 K. To check the effect of temperature, we ran the POPC system at 303 K. Again, the result was indistinguishable. The change occurred slightly more slowly than at 310 K.

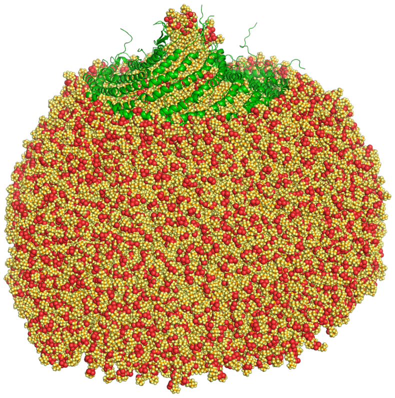

**Fig. S4**. 8S in POPC at 303 K.

e) All simulations used the charmm36m FF for the protein and c36 for the lipids. To check the effect of the force field, we ran the POPC system using Amber 19SB for the protein and Amber21 for the lipids. Very similar results were obtained.

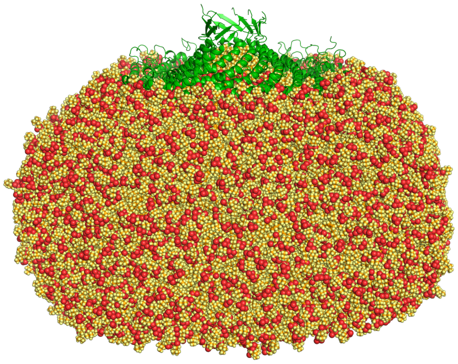

**Fig. S5**. 8S in POPC using Amber force fields.

f) Most simulations used the semi-isotropic barostat. In the membrane only system this resulted in a significant change in the aspect ratio of the unit cell. In contrast, the isotropic barostat ran without issues. To check the effect of the barostat on membrane curvature, several systems were also run with the isotropic barostat. The 9DN0 replicate is shown above. POPC and POPC/cholesterol are shown in Fig. S6. The results were very similar.

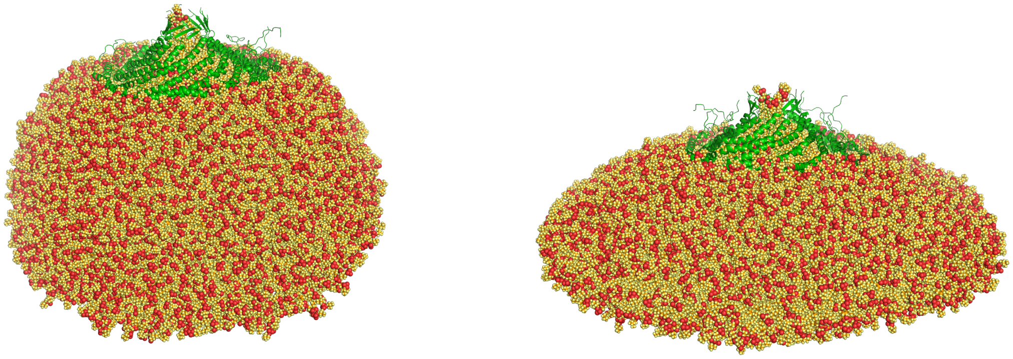

**Fig. S6**. 8S in POPC (left) and POPC/cholesterol (right) using the isotropic barostat.

g) To identify conditions under which the 8S complex remains flat, we ran a simulation in explicit n-hexane. The structure exhibited distortions, presumably due to the strong interactions between charged residues. The barrel half-dissolved, but did not rise as in the aqueous simulations.

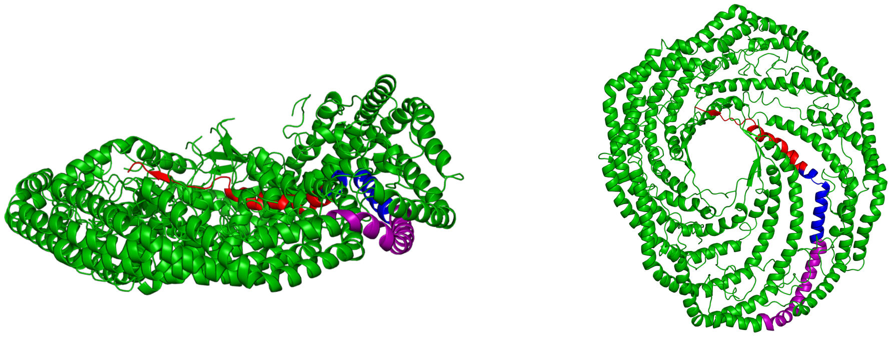

**Fig. S7**. Cav1 8S in explicit n-hexane, side and top views. One chain is colored according to domain.

h) The mutants P132L and S80E and cav2 are experimentally known to not form 8S complexes [1–3]. Nevertheless, we ran all three with the same protocol as cav1 and cav3 (a homology model using swissmodel was built for cav2). The P132L mutant gave very similar results to wild type (although in implicit solvent simulations, it remained flat). The S80E mutant also gave similar results to wild type. Cav2 remained flat and generated slightly negative curvature. Presumably, the 2 μs time scale is too short to reveal the thermodynamic instability of these complexes.

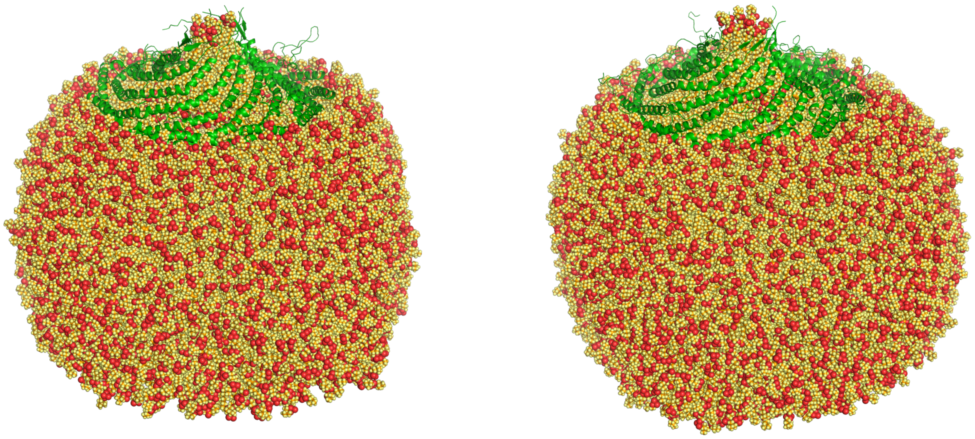

**Fig. S8**. P132L (left and S80E (right) in POPC.

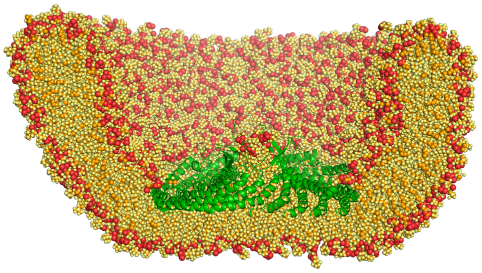

**Fig. S9**. Homology model of cav2 in POPC.

i) Cav1 palmitoylated at Cys133 and Cys156 was run in POPC and gave very similar results to the unpalmitoylated system. A model for full-length cav1 was constructed based on the AF2 prediction. That also gave very similar results to the truncated complex.

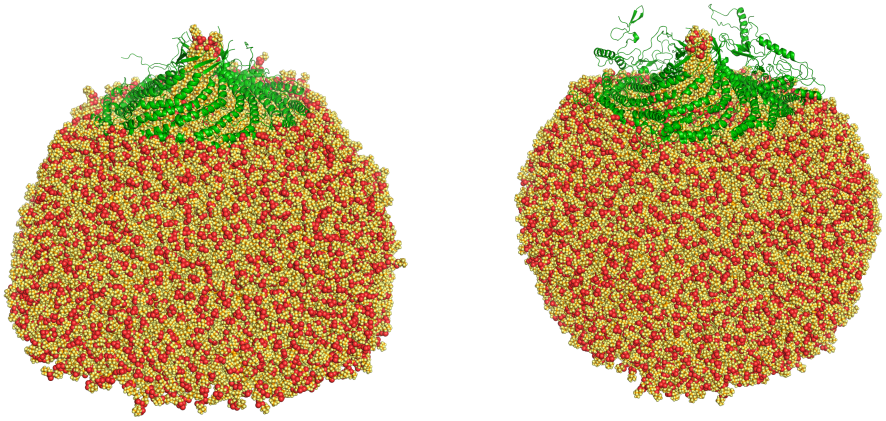

**Fig. S10**. Palmitoylated (left) and full-length cav1 (right) in POPC after 2 μs of simulation.
